# Structural basis of substrate recognition by human tRNA splicing endonuclease TSEN

**DOI:** 10.1101/2022.09.03.506465

**Authors:** Samoil Sekulovski, Lukas Sušac, Lukas S. Stelzl, Robert Tampé, Simon Trowitzsch

**Author notes:** Corresponding author: S.T.

## Abstract

The heterotetrameric human transfer RNA (tRNA) splicing endonuclease (TSEN) catalyzes the excision of intronic sequences from precursor tRNAs (pre-tRNAs)^1^. Mutations in TSEN and its associated RNA kinase CLP1 are linked to the neurodegenerative disease pontocerebellar hypoplasia (PCH)^2–8^. The three-dimensional (3D) assembly of TSEN/CLP1, the mechanism of substrate recognition, and the molecular details of PCH-associated mutations are not fully understood. Here, we present cryo-electron microscopy structures of human TSEN with intron-containing pre-tRNA^Tyr^gta and pre-tRNA^Arg^tct. TSEN exhibits broad structural homology to archaeal endonucleases^9^ but has evolved additional regulatory elements that are involved in handling and positioning substrate RNA. Essential catalytic residues of subunit TSEN34 are organized for the 3’ splice site which emerges from a bulge-helix configuration. The triple-nucleotide bulge at the intron/3’-exon boundary is stabilized by an arginine tweezer motif of TSEN2 and an interaction with the proximal minor groove of the helix. TSEN34 and TSEN54 define the 3’ splice site by holding the tRNA body in place. TSEN54 adapts a bipartite fold with a flexible central region required for CLP1 binding. PCH-associated mutations are located far from pre-tRNA binding interfaces explaining their negative impact on structural integrity of TSEN without abrogating its catalytic activity in vitro^10^. Our work defines the molecular framework of pre-tRNA recognition and cleavage by TSEN and provides a structural basis to better understand PCH in the future.

## Main

Transfer RNAs (tRNAs) rank among the most abundant RNA species in the cell playing a central role in protein biosynthesis and cellular homeostasis. tRNAs are transcribed as precursor molecules (pre-tRNAs), which undergo tightly controlled multistep processes for their maturation^11^. A subset of pre-tRNAs contains intronic sequences^12^, which are excised by the tRNA splicing endonuclease (TSEN)^1^, whereas the resulting 5’ and 3’ exons are ligated by the tRNA ligase complex^13^. Human TSEN comprises two catalytic subunits, TSEN2 and TSEN34, and two structural subunits, TSEN15 and TSEN54 (Fig. 1a), with an abundance of ~100 molecules per cell^1,14^. Little is known about the structure of human TSEN, in particular, how pre-tRNAs are recognized and how splice sites are defined. Only an NMR structure of homodimeric TSEN15 ^15^ and a crystal structure of a truncated TSEN15–34 heterodimer^10^ are available to date.

**Fig. 1.**
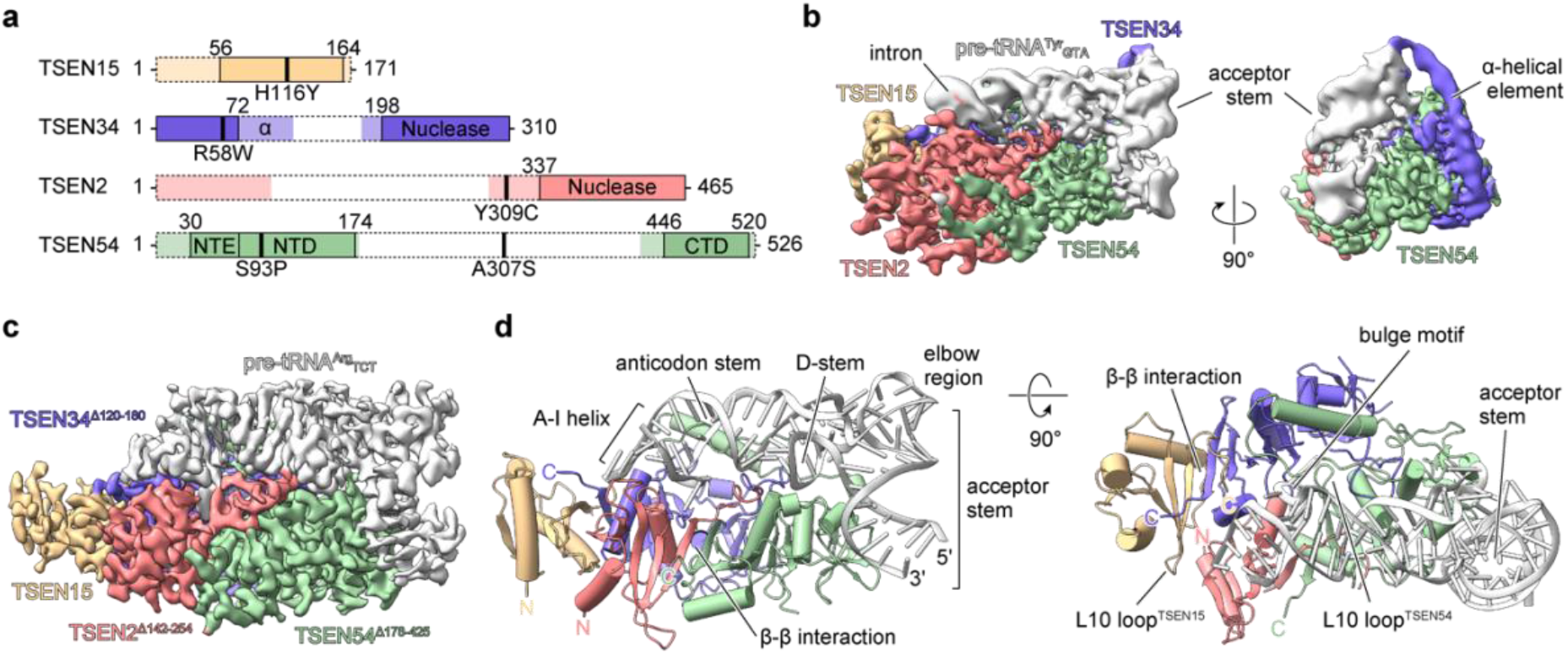
Structure of TSEN/pre-tRNA complexes. **a**, Bar diagrams of TSEN subunits depicting unstructured regions (dotted boxes) identified by cryo-EM analyses. Positions of pontocerebellar hypoplasia (PCH)-associated mutations are indicated. α – α-helical element, NTE – N-terminal extension, NTD – N-terminal domain, CTD – C-terminal domain. **b**, Cryo-EM density map (colored as in **a**) of full-length TSEN with pre-tRNA^Tyr^gta (grey) at 3.8 Å resolution. **c**, Cryo-EM density map of truncated TSEN with pre-tRNA^Arg^tct at 3.1 Å resolution. **d**, Corresponding atomic model (cartoon representation) of truncated TSEN/pre-tRNA^Arg^tct. N – amino terminus, C – carboxy terminus

Splicing endonucleases are classified into five families with different quaternary structures: α_4_, α’_2_, (αβ)_2_ and ε_2_ in Archaea^16,17^, and heterotetrametric αβγδ assemblies in eukaryotes^1,18^. In archaeal α_4_-type endonucleases, homotetramer formation is driven by the interaction of antiparallel β-sheets at the hydrophobic interface of two neighboring subunits and the ionic interaction between a negatively charged L10 loop and a positively charged pocket of an opposing subunit^9^. A crystal structure of truncated human TSEN15-34 confirmed these interactions in eukaryotes^10^. In humans, the predicted catalytic triad Tyr/His/Lys in TSEN2 and TSEN34 was shown to drive cleavage of the 5’ and 3’ splice site, respectively^10,19^.

tRNA introns are diverse in sequence and length^12^. At the intron-exon boundaries, archaeal pre-tRNAs form a central four base pair helix flanked by single-stranded three-nucleotide bulges, the bulge-helix-bulge (BHB) motif^20,21^. With the splice sites located within the bulges, the BHB motif was proposed as universal recognition element for archaeal intron cleavage at various positions in pre-tRNAs^22^. In eukaryotes, the intron position is strictly located at one nucleotide downstream of the anticodon^12,23^. This allows for a relaxed BHB motif with more variations in the central anticodon-intron (A-I) helix^23,24^. Only the proximal A-I base pair seems to be conserved among eukaryotes with a pivotal role for intron cleavage in yeast^23^, frogs^25,26^, fruit flies^27^, and humans^10,27^. In contrast to Archaea, eukaryotic splicing endonucleases are postulated to utilize TSEN54 (Sen54 in yeast) as a molecular ruler binding the mature pre-tRNA domain to define splice sites by distance^9,18,23,24,28^. The crystal structure of an archaeal endonuclease with bound BHB RNA revealed that two arginine residues act in trans at both active sites. By forming a cation-π sandwich, the residues pinch a bulge base to position the splice sites for cleavage^9,28,29^. Interestingly, analysis of yeast endonuclease verified the requirement of the cation-π sandwich only for 5’ cleavage but not for 3’ cleavage^29^.

Mammalian TSEN additionally associates with the RNA kinase CLP1 by uncharacterized interactions^1,30^. While in vitro experiments showed that CLP1 is dispensable for pre-tRNA splicing^10,19^, in vivo and in-cell studies connected mutations in CLP1 to impaired pre-tRNA splicing and neurodegeneration^7,8,31–33^. Indeed, mutations in CLP1 and all four subunits of human TSEN have been associated with pontocerebellar hypoplasia (PCH), a broad group of inherited neurodegenerative disorders^2–8,31,34^.

Here we purified human TSEN and determined the single-particle cryo-EM structures in complex with intron-containing pre-tRNA^Tyr^gta and pre-tRNA^Arg^tct. These structures define key factors for substrate recognition and intron excision and localize the PCH-associated mutations in their molecular environment. We provide evidence that CLP1 is flexibly associated to TSEN via an unstructured region of TSEN54 supporting its proposed function as a negative regulator of tRNA splicing in humans^19,35,36^.

## Structure determination and overall architecture of TSEN

Biochemical studies had previously revealed that the four TSEN subunits TSEN2, TSEN15, TSEN34, and TSEN54 are necessary and sufficient for cleavage at the 5’ and 3’ splice sites of human precursor tRNAs^10,19^. To better understand pre-tRNA recognition by human TSEN, we purified recombinant, full-length TSEN which was rendered catalytically inactive by introduction of the active site mutants TSEN2^H377F^ and TSEN34^H255F 10^. Inactive TSEN was complexed with pre-tRNA^Tyr^gta (Extended Data Fig. 1), which is the most abundant intron-containing tRNA in humans and whose isodecoders are only present as intron-containing genes^12,37^. Monodisperse TSEN/pre-tRNA^Tyr^gta complexes were isolated by size exclusion chromatography and vitrified on cryo-EM grids (Extended Data Figs. 1,2). The 3.8-Å cryo-EM reconstruction revealed a compact particle that readily displayed the relative orientation of the pre-tRNA and protein subunits (Fig. 1b). The map quality allowed fitting of predicted 3D structures for all TSEN subunits^38^, but we could not unambiguously build a complete atomic model due to variations in local map resolution (Extended Data Fig. 3a). However, the cryo-EM reconstruction revealed a predicted extended α-helical element in TSEN34 that may assist in positional proofreading of pre-tRNAs by sensing their elbow region (Fig. 1b). Large parts of TSEN polypeptide chains and the tip of the intron were not visible in the cryo-EM map (Fig. 1a,b). We reasoned that these unstructured/flexible regions impeded high resolution reconstructions. Consequently, we truncated unresolved central regions of TSEN subunits and utilized pre-tRNA^Arg^tct which exhibits the shortest intronic sequence of intron-containing pre-tRNAs^12^. Indeed, the cryo-EM reconstruction process and the final map quality significantly profited from using a truncated TSEN complex and pre-tRNA^Arg^tct (Fig. 1c,d, Extended Data Fig. 2e–h, and Extended Data Fig. 3b). The 3.1-Å cryo-EM reconstruction of the truncated ribonucleoprotein complex revealed an identical architecture as seen for full-length TSEN with pre-tRNA^Tyr^gta (Fig. 1b,c) and enabled us to unambiguously build a model with good stereochemistry (Fig. 1d, Extended Data Fig. 3c–g, and Extended Data Table 1).

## Subunit–substrate interactions within human TSEN

The two structural subunits TSEN15 and TSEN54 pair with their catalytically active counterparts TSEN34 and TSEN2 in a tail-to-tail fashion via their C-terminal β-strands (Fig. 2). All TSEN subunits contain a characteristic C-terminal nuclease fold (C-terminal domain, CTD) that is built up by a central, twisted, 5-stranded β-sheet sandwiched by two α-helices (Fig. 2a). Whereas TSEN2 and TSEN15 display flexible N-terminal regions, TSEN34 and TSEN54 contain large unstructured central domains of different length (Fig. 1a). The N-terminal domains (NTDs) of TSEN34 and TSEN54 both fold into a barrel-like, 5-stranded β-sheet structure stabilized by a short α-helix (Fig. 1d, Fig. 2a, Extended Data Fig. 4a). Whereas polar contacts dominate the binding interface of opposing subunits in archaeal tRNA endonucleases including the acidic L10 loop, TSEN34 and TSEN54 engage each other by an intricate interface involving hydrogen bonds, salt bridges, and hydrophobic interactions with a contact surface area of 2287.2 A^2^ (Fig. 2b). An N-terminal extension (NTE) of TSEN54 clamps a loop structure of the NTD of TSEN34 against the NTD of TSEN54 leading to a tight, interlocked subunit coalescence (Fig. 2b). This configuration allows TSEN34 and TSEN54 to assist each other in positioning the mature domain of pre-tRNAs by polar contacts with the phosphoribose backbone of the acceptor stem, the D-stem, and the anticodon stem (Fig. 2c–e and Extended Data Fig. 5). The extensive TSEN34–54 contact surface to the pre-tRNA covers an area of 1838.3 A^2^, explaining why mature tRNAs are also bound by TSEN with similar dissociation constants as pre-tRNAs^10^. All-atom molecular dynamics (MD) simulations (1 μs time scale) show a highly rigid core structure of the complex with flexibility only at the periphery including the tip of the intron, the elbow region, the 5’ and 3’ ends, and the central domain of TSEN2 (Extended Data Fig. 1d). Whereas no interaction between pre-tRNA and TSEN15 was evident from our structure, we found TSEN2 to contact the G46-U34 wobble base pair of the A-I helix. By formation of a hydrogen bond between the side chain of N413 and the N^2^-amine of G46 (Extended Data Fig. 3g), TSEN2 may be involved in sensing A-I helix stability^27^.

**Fig. 2.**
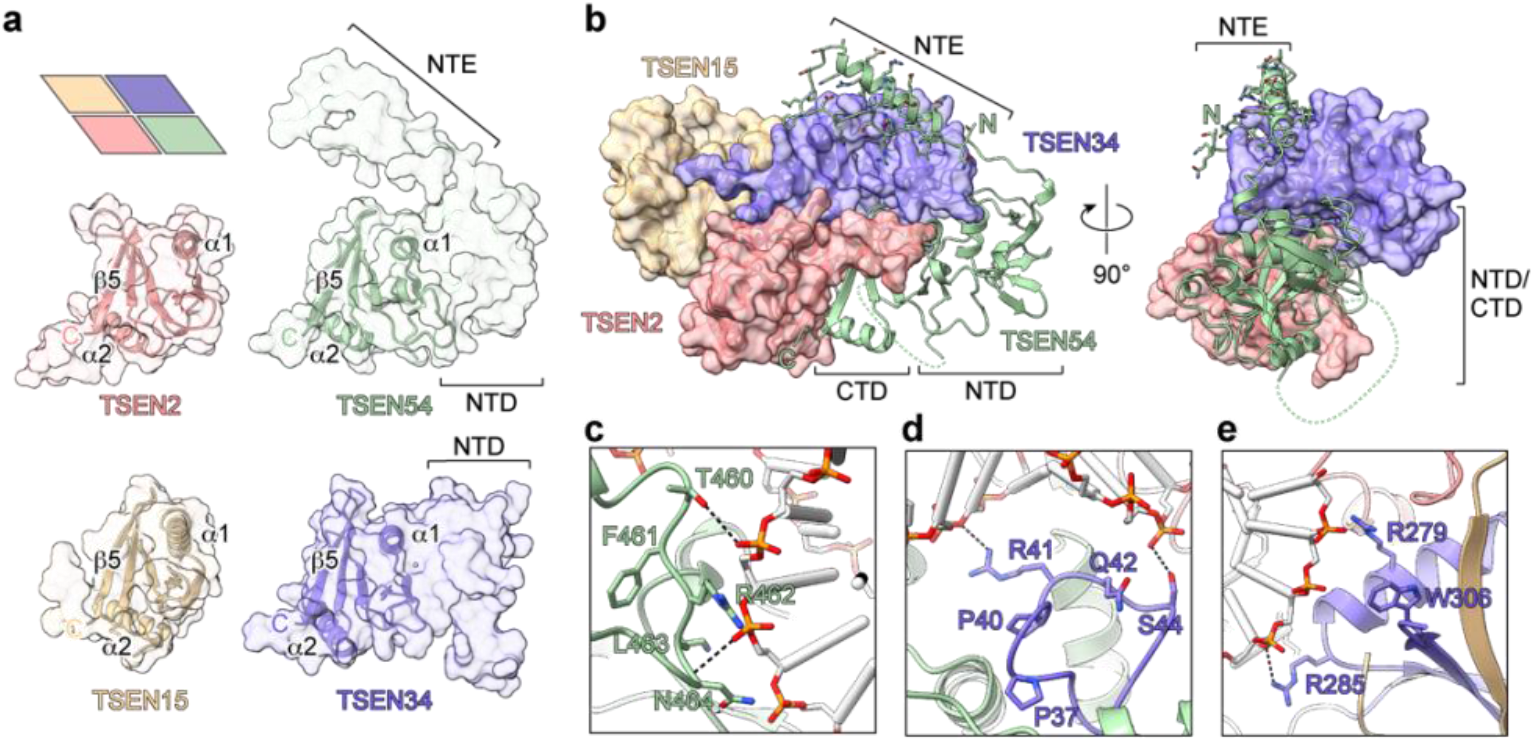
Interaction networks between TSEN subunits and pre-tRNA substrate. **a**, Surface representation of individual TSEN subunits highlight common folds of their C-terminal domains (CTD, cartoon representation). CTDs are oriented to match the TSEN54 view in panel **b**. Inset shows schematic illustration of intersubunit orientations as observed in the assembled complex. Carboxy termini (C) are indicated. **b**, Interface of TSEN54 (pale green cartoon model) with TSEN2-15-34 (colored cartoon-surface model). Pre-tRNA is omitted for clarity. NTE – N-terminal extension, NTD – N-terminal domain, CTD – C-terminal domain, N – amino terminus. **c-e**, Close-up views of TSEN/pre-tRNA^Arg^tct interactions. Key residues are highlighted. **c**, Acceptor stem phosphates (yellow and red) make polar contacts with TSEN54 loop (green). **d** Polar interactions of TSEN34 (slate blue) with D-stem phosphate backbone. **e**, TSEN34-anticodon-stem interface in vicinity to 5’ splice site.

## Molecular details governing the 3’ splice site selection

Outside of the intronic region, pre-tRNA^Arg^tct adopts a near identical conformation as a matured tRNA evidenced by structural superposition to *E. coli* tRNA^Phe^gaa ^39^ with a root mean square deviation (RMSD) of 2.7 Å (Extended Data Fig. 4b). The pre-tRNA’s anticodon stem forms an A-form helix which extends from the A31-U52 base pair into a 4-base pair A-I helix initiated by the A-I base pair (C32-G48) (Fig. 3a,b). The last bases of the A-I helix form a non Watson–Crick pair (Fig. 3a) explaining the observed requirement of several weak base pairs within the A-I helix for efficient tRNA splicing^27^. The 3-nucleotide bulge motif (A49-A51) harboring the 3’ splice site forms a knot-like structure that expels the nucleobases from the A-form helix for interactions with TSEN proteins (Fig. 3b). Reminiscent to the architecture of the 3’ splice site in archaeal endonucleases^28^, the nucleobase of the first bulge nucleotide, A49, is sandwiched by R409 and R452 of TSEN2 forming a well-conserved cation-π stack (Fig. 3d). The guanidinium group of R409 of TSEN2 hydrogen bonds with the 2’-hydroxyl oxygen of A49 and the phosphate of U52 stabilizing the knot-like structure of the bulge (Fig. 3d). Furthermore, K239 of TSEN34 additionally locks the conformation of the pre-tRNA backbone by contacting the phosphate of A50 (Fig. 3d). In addition to the formation of the cation-π sandwich with TSEN2, the tight looping of the bulge brings A51 into vicinity of the A-I helix’ minor groove and allows a base-to-ribose hydrogen bond between the adenine ring of A51 and the 2’-hydroxyl oxygen of C32 (Fig. 3b,c). In the structure of archaeal endonuclease with BHB RNA, the corresponding adenine forms an additional hydrogen bond with a G nucleobase thereby resulting in an obligate C-G pairing at the A-I base pair position in Archaea^25,26,28,40^. Our structure shows that human TSEN only relies on the presence of any A–I base pair in a sequence independent manner for cleavage at the 3’ splice site (Fig. 3b,c), explaining recent biochemical data^10,27^.

**Fig. 3.**
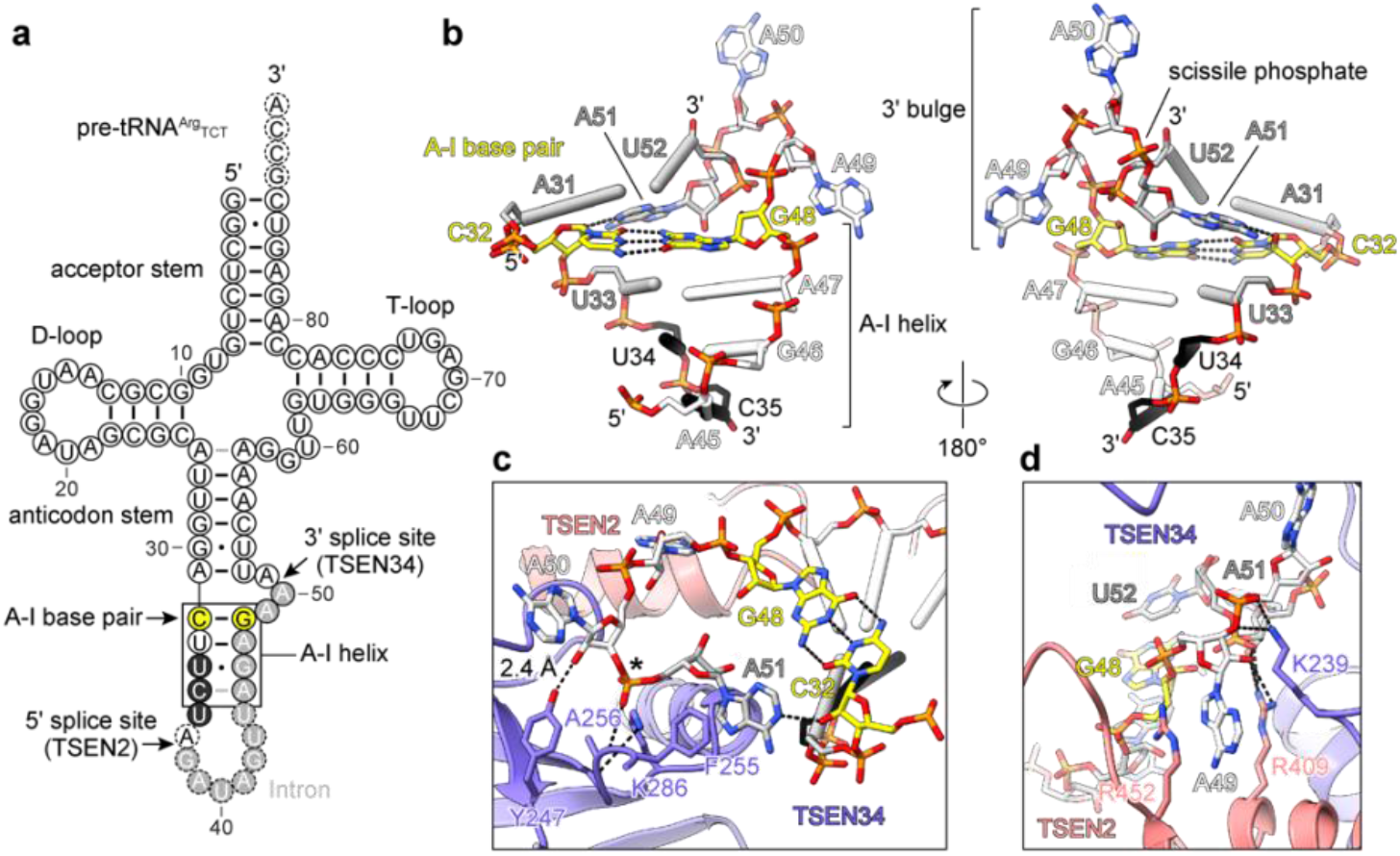
Molecular determinants of 3’ splice site selection. **a**, Secondary structure cloverleaf model of pre-tRNA^Arg^tct. Intron nucleotides are shown as white letters and grey background. Arrows indicate intronic splice sites and the anticodon-intron (A-I) base pair (yellow) as part of the A-I helix (box). Dashed circles mark nucleotides not resolved in the cryo-EM model. **b**, Structure of the A-I helix. The hydrogen bonding pattern of 3’ bulge base A51 and A-I base pair C32-G48 are highlighted. **c**, Magnified view of the 3’ splice site. The TSEN34 (slate blue) catalytic triad consisting of Y247, H255 (mutated to F in inactive TSEN) and K286 is shown as stick models. Polar contacts (dashed lines) of A256 peptide backbone with scissile phosphate (asterisk) and stacking of F255 and A51 show interaction network prior to cleavage. **d**, Detailed structure of 3’ bulge cation-π-sandwich. TSEN2 (salmon) residues R409 and R452 together with TSEN34 (slate blue) K239 interact with A49 to form the composite active splice site.

Mutagenesis studies in both archaeal and yeast tRNA-splicing endonucleases have identified a conserved histidine residue as part of a catalytic triad critical for cleavage catalysis^9,41^. Opposite the TSEN arginine tweezer pair, residues Y247, H255, and K286 of TSEN34 are found in the same configuration as their yeast and archaeal counterparts^18,28,29^ (Fig. 3c). The scissile phosphate is locked into position by a hydrogen bond to the backbone nitrogen of A256. The side chain oxygen of Y247 is optimally positioned for activation of the 2’-hydroxyl oxygen of A50 consistent with the proposed mechanism which postulates that Y247 deprotonates the nucleophilic 2’-oxygen, H255 protonates the leaving 5’ hydroxyl oxygen and K286 stabilizes the developing negative charge of the transition state^28^. A structurally conserved 3’ splice site in TSEN agrees with data from Xenopus splicing endonuclease, which was shown to cleave a miniprecursor tRNA containing only the mature domain and the 3’ splice site^26^.

Previous biochemical work on the yeast splicing endonuclease had concluded that 3’ splicing was independent of the cation-π sandwich of Sen2 and only depended on the conserved A-I base pair^29^. Counter to this postulated mechanism, we observe cleavage at the 3’ splice site resembles the requirements of a bulge-helix motif including the cation-π sandwich, brought about by TSEN2, as well as the A-I helix and base pair (Fig. 3). The cation-π sandwich thought to be required for the 5’ splice site is derived from TSEN34, which provides the catalytic triad at the 3’ splice site. Indeed, we find R279 and W306 of TSEN34 at the corresponding positions as the arginine tweezer pair (R280 and R302) of *Archaeoglobus fulgidus* endonuclease^28^, albeit not involved in pinching out a nucleotide base (Fig. 2e). Whereas the scissile phosphate at the 3’ splice site is readily positioned towards the active site residues of TSEN34, cleavage at the 5’ splice site necessitates structural rearrangements and presumably melting of the A-I helix. This implies that there must be differences between cleavage rates at the 3’ and 5’ splice sites, and potentially an ordered sequence of events. A recent bioinformatic analysis of RNA ends in yeast identified novel messenger RNA (mRNA) substrates of TSEN, which share a conserved adenosine at the −1 position of the 3’ cleavage site^42^. Since all mRNA substrates were shown to be dependent on the catalytic activity of TSEN2, but not on TSEN34^42^, we speculate that the high structural plasticity of TSEN at the 5’ splice site allows the enzyme to recognize substrates other than pre-tRNAs.

## Insights into pontocerebellar hypoplasia-associated disease mutations

Mutations in TSEN^2–6,34^ and CLP1^7,8^ are associated with pontocerebellar hypoplasia (PCH) in humans. PCH collectively describes a heterogeneous group of early onset neurodegenerative disorders with common symptoms as underdevelopment of the pons and cerebellum and concomitant microcephaly^34^. We previously demonstrated a destabilization of mutated TSEN causing insufficient tRNA splicing events in PCH patient-derived fibroblasts while catalytic activity in biochemical assays was unaffected^10^. The TSEN/pre-tRNA^Arg^tct structure, determined in the present work, now allowed us to localize and rationalize the effects of TSEN mutations in the context of the pre-tRNA-engaging multiprotein complex (Fig. 4a). The S93P mutation located in a loop in vicinity of the pre-tRNA acceptor stem would introduce additional conformational rigidity with P94, putatively disturbing the hydrophobic interface between the adjacent α-helix and β-sheets (exemplarily indicated by L91 and F99) (Fig. 4b). At the TSEN15–34 interface, TSEN15 H116 is the only known PCH-associated residue mediating interaction between two subunits (Fig. 4c). Mutation H116Y is proposed to interrupt hydrogen bonding and cause steric clashes at the interface. Similarly, TSEN34 R58 engages in a hydrogen bonding network which would be abolished by introduction of the bulky tryptophan mutation (R58W) but also introduce large steric clashes (Fig. 4d). The severity of the postulated effects correlates well with the previously reported decrease in denaturing temperature of mutated TSEN complexes (S93P: T_Δ_ = −5.1 °C, H116Y: T_Δ_ = −3.0 °C, R58W: T_Δ_ = −6.2 °C)^10^. Based on our structural analysis, we propose that the rare PCH-associated mutations TSEN15 W76G, TSEN15 Y152C, TSEN54 E85V, and TSEN54 Y119D lead to similar thermal destabilization of TSEN (Extended Data Fig. 6). Overall, these data illustrate how PCH-associated mutations distant from pre-tRNA binding surfaces negatively impact structural integrity in vivo but not catalytic activity in vitro^10^.

**Fig. 4.**
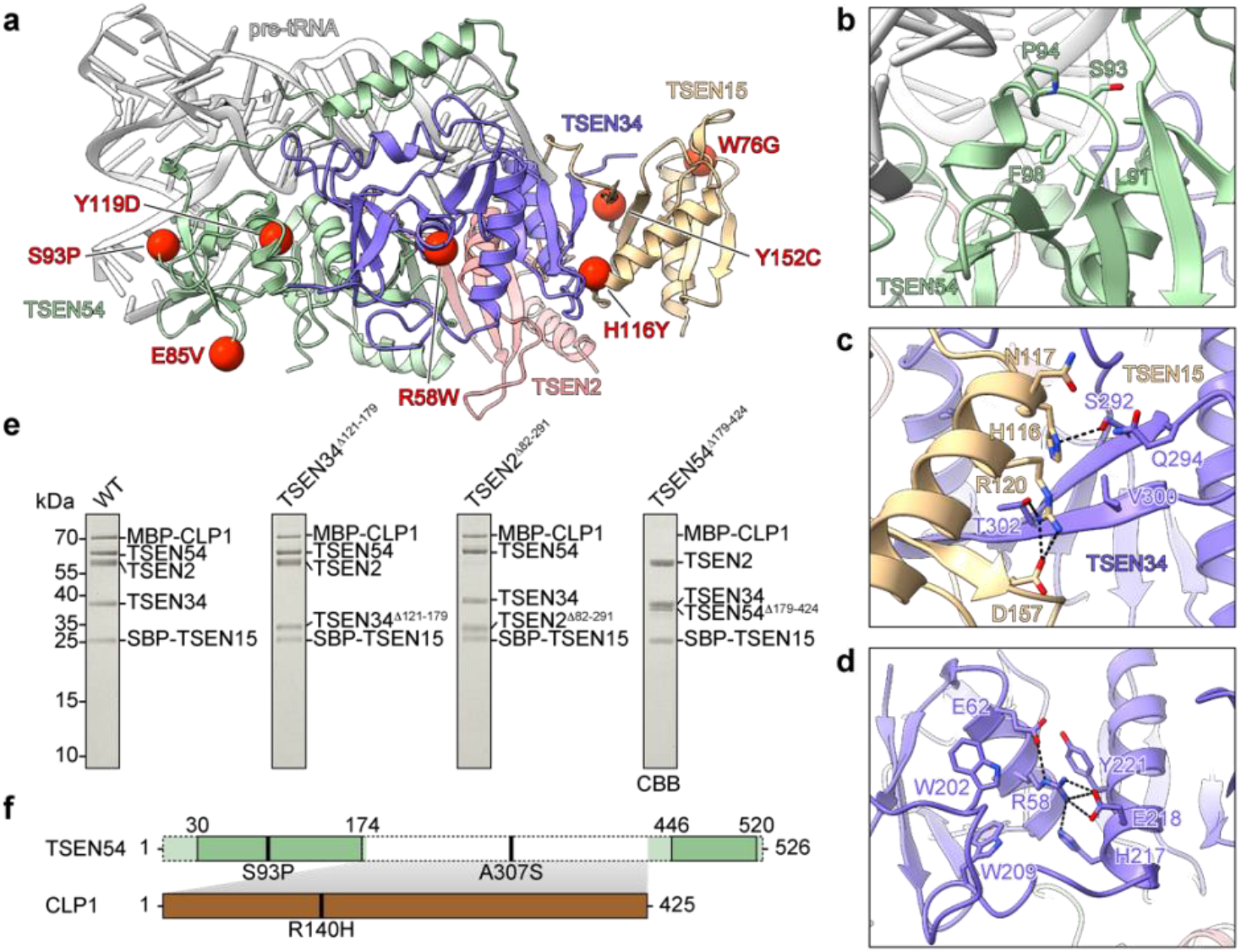
Structure-guided assessment of PCH-associated mutations and effects on CLP1 binding. **a**, Global positions of PCH-associated mutations (red spheres) in TSEN subunits TSEN15 (yellow), TSEN34 (slate blue), and TSEN54 (green). **b-d**, Molecular details of TSEN interactions in regions of disease mutation hotspots. **b**, TSEN54 S93 environment (green) in proximity to pre-tRNA acceptor stem (grey). **c**, TSEN15 (yellow) H116 at binding interface with TSEN34 (slate blue). **d**, TSEN34 R58 environment. **e**, Effect of deletion of disordered TSEN linker regions on MBP-CLP1 binding. SBP-pulldowns were visualized by SDS-PAGE. Descriptions above gels indicate substitutions of TSEN subunits. WT – wild-type TSEN/MBP-CLP1. Gels are representative of n = 3 biologically independent experiments. **f**, Bar diagram of proposed TSEN54-CLP1 interaction (grey-shaded). The TSEN54 unstructured region (dotted white box) putatively interacts with CLP1. Positions of PCH-associated mutations are mapped with black strokes.

The RNA kinase CLP1, a part of the pre-mRNA cleavage factor IIm in humans^43^, was shown to associate with TSEN and to phosphorylate the 5’ end of the 3’ tRNA exon^1,30^. However, the role of CLP1 during tRNA splicing and the assembly of CLP1 with TSEN remains enigmatic. Based on our structure we hypothesized that a disordered region of TSEN recruits CLP1, similar to the association of yeast Clp1 to CFIA via Pcf11 in yeast^44^. We performed pull-down experiments of upon overexpression of TSEN/CLP1 complexes harboring single substitutions of truncated TSEN subunits (Fig. 4e). These assays revealed that deletion of the disordered central region of TSEN54 abrogates CLP1 binding while the other substitutions do not impair CLP1 binding (Fig. 4e). These data indicate the TSEN54 linker region (residues 179 to 424) as interaction hub for CLP1 recruitment to TSEN (Fig. 4f). Strikingly, the most commonly found PCH-associated mutation TSEN54 A307S maps to the middle of this linker region suggesting a role in CLP1 recruitment (Fig. 4f).

## Summary

Our structure of human TSEN in complex with pre-tRNA substrate reveals long-sought molecular details for complex assembly and cleavage catalysis. For the first time, allowing comparisons of human TSEN with its archaeal endonuclease counterparts^28^ which highlight common and divergent features in splicing endonuclease evolution. Crucial molecular details of substrate handling are unveiled, revealing a structural asymmetry in formation of the 5’ and 3’ splice sites. Since the order of cleavage events at the 5’ and 3’ splice sites of yeast pre-tRNAs appears random^45^, structures of TSEN at different stages of catalysis will be instrumental for a comprehensive description of the cleavage pathway in yeast and humans. Our structural data explain why the scissile phosphate at the 3’ splice site invariably precedes residue 38 of the mature tRNA, demonstrating that TSEN54 indeed acts as a molecular ruler. Finally, we provide a structural basis for a deeper understanding of disease mutations leading to TSEN-associated neurological disorders.

## Methods

No statistical methods were used to predetermine sample size. The experiments were not randomized, and investigators were not blinded to allocation during experiments and outcome assessment.

### Plasmid constructs

Open reading frames encoding the TSEN subunits TSEN2 (UniProtKB Q8NCE0), TSEN15 (UniProtKB Q8WW01), TSEN34 (UniProtKB Q9BSV6), TSEN54 (UniProtKB Q7Z6J9), and CLP1 (UniProtKB Q92989) were amplified by polymerase chain reaction (PCR) and cloned into the modified MultiBac vectors pAMI, pMIDC, pMIDK, and pMIDS as described ^10^. An N-terminal streptavidin-binding peptide (SBP) tag or a His10-tag followed by a Tobacco Etch Virus (TEV) protease cleavage site were engineered for TSEN15 by restriction cloning. The active site mutations in TSEN2 (TSEN2^H377F^) and TSEN34 (TSEN34^H255F^) were introduced via QuikChange mutagenesis. The N-terminal fusion of maltose binding protein (MBP) to CLP1 was generated via overlap extension PCR. A 250-bp gene segment of CLP1 was extended with a linearized MBP gene fragment containing a glycine-serine linker and a TEV protease cleavage site. The fragment was used to replace a short CLP1 sequence in frame in the plasmid encoding CLP1. This procedure yielded an MBP-TEV-CLP1 construct. Truncated TSEN subunits TSEN2^Δ82-291^, TSEN34^Δ121-179^ and TSEN54^Δ179-424^ were generated by deletion of bona fide disordered regions according to the full-length TSEN cryo-EM model. To this end, the sequences encoding the disordered regions in the respective plasmids were replaced by glycine-serine linkers using sequence and ligation independent cloning (SLIC)^46^. For generation of a full-length TSEN encoding bacmid, respective acceptor and donor vectors were concatenated by Cre-mediated recombination utilizing the LoxP sites present on each vector. Consequently, the fusion plasmid was integrated into the EMBacY baculoviral genome via Tn7 transposition.

Human genes encoding pre-tRNA^Tyr^gta8-1 and pre-tRNA^Arg^tct3-2 and the yeast gene encoding pre-tRNA^Phe^gaa2-2 were amplified by PCR from genomic DNA of human embryonic kidney (HEK293) cells and *Saccharomyces cerevisiae* strain S288C, respectively. Pre-tRNA sequences were flanked by a preceding T7 promoter sequence and pre-tRNA^Tyr^gta8-1 was additionally optimized for in vitro transcription by placing a GG at the starting position and a CC at pairing positions in the acceptor stem. DNA fragments were cloned into a pUC19 vector via sticky end ligation using *Bam*HI and *Hind*III restriction sites. All constructs were verified by Sanger sequencing. A list of all primers used in this study is provided in Supplementary Table 1.

### Production and purification of human TSEN complexes

Recombinant truncated human TSEN complexes were produced in human embryonic kidney (HEK) 293F cells by cotransfection using linear polyethyleneimine (PEI) (MW 25000, Polysciences) and incubated for 72 h at 37 °C. Typically, TSEN complexes were produced in 0.5 liters of HEK293F cell suspension culture at a cell density of 1 × 10^6^ cells ml^−1^ by transfection with 750 μg of plasmid DNA. Cells were harvested by centrifugation at 500 × g for 10 min. Cell pellets were flash-frozen in liquid nitrogen and stored at −80 °C until further use. Cell pellets were resuspended in 10 ml of lysis buffer comprising 50 mM HEPES-NaOH, pH 7.4, 400 mM NaCl, 0.5 mM phenylmethanesulfonyl fluoride (PMSF), 1.25 mM benzamidine, per 100 ml expression volume and lysed by sonication. Lysates were cleared by centrifugation at 50,000 × g for 40 min in a Type 45 Ti fixed-angle rotor (Beckman Coulter). Pre-equilibrated Pierce High Capacity Streptavidin agarose resin (Thermo Fisher Scientific) was added to the soluble fraction and incubated for 1 h at 4 °C under agitation. The resin was recovered by centrifugation and washed extensively in lysis buffer without protease inhibitors. Bound proteins were eluted in 50 mM HEPES-NaOH, pH 7.4, 400 mM NaCl, 2.5 mM biotin. Eluates were subjected to TEV protease cleavage (1:20 protease to protein mass ratio) at 4 °C to remove the SBP-tag, concentrated by ultrafiltration using Amicon Ultra centrifugal filters (Merck) with a molecular weight cut-off (MWCO) of 30 kDa and polished by size exclusion chromatography on a Superdex 200 Increase 3.2/300 column (GE Healthcare) equilibrated in 50 mM HEPES-NaOH, pH 7.4, 400 mM NaCl. Peak fractions were pooled, concentrated by ultrafiltration, and flash-frozen in liquid nitrogen after supplementation with 15% (v/v) glycerol.

Production of baculoviruses and overproduction of recombinant human TSEN complexes in *Spodoptera frugiperda* (*Sf*) 21 cells was performed as described^10^. Typically, TSEN complexes were produced in 1.6 liters of *Sf*21 suspension culture at a cell density of 1 × 10^6^ cells ml^−1^ by infection with 0.5–1% (v/v) of V1 baculovirus supernatant. 72 h post cell proliferation arrest, insect cells were harvested by centrifugation at 800 × g for 5 min. Cell pellets were flash-frozen in liquid nitrogen and stored at −80 °C until further use.

Insect cell pellets were resuspended in 10 ml of lysis buffer comprising 50 mM HEPES-NaOH, pH 7.4, 400 mM NaCl, 10 mM imidazole, 0.5 mM phenylmethanesulfonyl fluoride (PMSF), and 1.25 mM benzamidine, per 100 ml expression volume, and lysed by sonication. Lysates were cleared by centrifugation at 50,000 × g for 40 min in a Type 45 Ti fixed-angle rotor (Beckman Coulter). Preequilibrated Ni^2+^-nitrilotriacetic acid (NTA) agarose resin (Thermo Fisher Scientific) was added to the soluble fraction and incubated for 45 min at 4 °C under agitation. Agarose resin was recovered by centrifugation and washed extensively in lysis buffer without protease inhibitors. Bound proteins were eluted in 50 mM HEPES-NaOH, pH 7.4, 400 mM NaCl, 250 mM imidazole. Eluates of immobilized metal ion affinity chromatography (IMAC) were subjected to TEV protease cleavage (1:20 protease to protein mass ratio) at 4 °C to remove the His-tag, concentrated by ultrafiltration using Amicon Ultra centrifugal filters (Merck) with a molecular weight cut-off (MWCO) of 30 kDa and polished by size exclusion chromatography on a Superdex 200 Increase 10/300 GL column (GE Healthcare) equilibrated in 50 mM HEPES-NaOH, pH 7.4, 400 mM NaCl. Peak fractions were pooled, concentrated by ultrafiltration, and flash-frozen in liquid nitrogen after supplementation with 15% (v/v) glycerol.

### Pull-down assays

For pulldown assays of SBP-tagged TSEN with MBP-CLP1 or TSEN substituted with the respective truncated subunit (TSEN2^Δ82-291^, TSEN34^Δ121-179^, or TSEN54^Δ179-424^) and MBP-CLP1, adherent HEK293T cells were transfected with the expression plasmids using linear PEI (MW 25000, Polysciences). In detail, 9 × 10^6^ HEK293T cells were seeded the day before transfection in 150 mm dishes in Dulbecco’s modified Eagle medium (DMEM, Gibco Life Technologies) with 10% fetal bovine serum (FBS, Capricorn Scientific) and incubated at 37 °C, 5% CO_2_, and 90% humidity. After 24 h, cells were transfected with 30 μg of DNA and a 1:3 ratio of PEI per 150 mm dish. Transfected cells were further incubated for 72 h, detached by addition of Dulbecco’s phosphate buffered saline (DPBS, Thermo Fisher Scientific) and harvested by centrifugation at 500 x g for 10 min. The cell pellets were flash frozen in liquid nitrogen and stored at −80 °C until further use. Frozen cell pellets were thawed and resuspended in 3 ml of lysis buffer containing 50 mM HEPES-NaOH, pH 7.4, 400 mM NaCl, 0.5 mM PMSF, 1.25 mM benzamidine, per 150 mm dish and lysed by sonication. Lysates were cleared by centrifugation at 20,817 × g for 1 h. Pre-equilibrated Pierce High Capacity Streptavidin agarose resin (Thermo Fisher Scientific) was added to the soluble fraction and incubated for 1 h at 4 °C under agitation. Agarose resin was recovered by centrifugation and washed extensively in lysis buffer without protease inhibitors. Bound proteins were eluted in 50 mM HEPES-NaOH, pH 7.4, 400 mM NaCl, 2.5 mM biotin. Eluates were mixed with 4x SDS sample buffer, incubated at 95 °C for 5 min and analyzed by SDS-PAGE and subsequent InstantBlue Coomassie (Abcam) staining.

### Run-off in vitro transcription and purification of pre-tRNAs

Genes encoding *S.c*. pre-tRNA^Phe^GAA2-2 and human pre-tRNA^Tyr^gta8-1 and tRNA^Arg^tct3-2 were amplified by PCR of respective pUC19 vectors and template DNA was isolated using the NucleoSpin Gel and PCR clean-up kit (Macherey-Nagel). RNA substrates were produced by run-off in vitro transcription using T7 RNA polymerase (New England Biolabs) and purified via anion exchange chromatography as described before with slight modifications ^47,48^. Briefly, 1 μg ml^−1^ of template DNA was mixed with 1000 U ml^−1^ of T7 polymerase and 1.5 mM of each rNTP (New England Biolabs) in 40 mM Tris-HCl, pH 7.9, 6-9 mM MgCl_2_, 2 mM spermidine, 1 mM DTT, and incubated for 4 h at 37 °C. To isolate transcribed RNAs, the reaction mixture was diluted in a 1:1 ratio (v/v) with AEX buffer comprising 50 mM sodium phosphate, pH 6.5, 0.2 mM EDTA, and loaded onto a HiTrap DEAE FF column (GE Healthcare) equilibrated in AEX buffer and eluted by a linear gradient from 0 to 700 mM NaCl. RNA containing fractions were analyzed via denaturing urea-PAGE with subsequent toluidine blue staining. RNAs were concentrated by ultrafiltration using Amicon Ultra 3 MWCO centrifugal filters (Merck) and stored at −80 °C. 10 pmol TSEN complexes were mixed with the respective RNA in a 1:2.5 molar ratio in 50 mM HEPES-NaOH, pH 7.4, 100 mM NaCl, 4 mM MgCl_2_, 1 mM DTT in a 20 μl reaction volume and incubated at 37 °C for 1.5 h. Reactions were stopped by the addition of a 2× RNA loading buffer (95% formamide, 0.02% SDS, 1 mM EDTA) and incubation at 70 °C for 10 min. Reaction products were separated by denaturing urea-PAGE and visualized by toluidine blue staining.

### Cryo-EM data collection and image processing

Purified, inactive human TSEN carrying the active site mutants TSEN2^H377F^ and TSEN34^H255F^ (18 μM) was incubated with pre-tRNA^Tyr^gta8-1 at 27 μM for formation of the protein–RNA complex. The complex was purified using a Superdex 200 Increase 3.2/300 gel filtration column (GE Healthcare) in a buffer comprising 10 mM HEPES-NaOH, pH 7.4, 200 mM NaCl, 1.5 mM MgCl_2_. Peak fractions were collected and 3 μl of TSEN/pre-tRNA complex were applied onto freshly glow-discharged copper grids (Quantifoil, Cu R1.2/1.3) and plunge-frozen in liquid ethane using a Vitrobot Mark IV (Thermo Fisher Scientific). Micrographs of full-length TSEN with pre-tRNA^Tyr^gta were recorded automatically (EPU) on a 300-kV FEI Titan Krios cryo-transmission electron microscope (cryo-TEM) in electron counting mode with a Falcon 3EC direct electron detector (Thermo Scientific) at a nominal magnification of ×75,000 corresponding to a calibrated pixel size of 1.06 Å. Dose-fractionated movies were acquired at an electron flux of 1.22 e^−^ per pixel per s over 75.0 s with 1.6 s exposures per frame (47 frames in total), corresponding to a total electron dose of ~81 e^−^ Å^2^. Movies were recorded in the defocus range from −1.0 to –2.5 μm (Extended Data Table 1).

Purified inactive truncated human TSEN carrying the TSEN2^H377F^ and TSEN34^H255F^ mutations (18 μM) was incubated with pre-tRNA^Arg^tct3-2 at 27 μM for formation of the protein–RNA complex. Purification and application to grids was performed as described above. Micrographs were recorded automatically (EPU) on a 200-kV Glacios cryo-TEM (Thermo Scientific) equipped with a Falcon 3EC direct electron detector (Thermo Fischer). The nominal magnification was ×190,000 resulting in a pixel size of 0.78 Å. Dose-fractionated movies were acquired at an electron flux of 0.8 e^−^ per pixel per s over 44.5 s with 0.9 s exposures per frame (48 frames in total), corresponding to a total electron dose of ~63 e^−^ Å^2^. Movies were recorded in the defocus range from −1.0 to –2.5 μm (Extended Data Table 1).

All image processing was done in cryoSPARC v3.0.1 - 3.3.1 (Structura Biotechnology Inc.)^49^. For full-length TSEN-pre-tRNA^Tyr^gta, a total of 2,231 movies from two datasets were used for analysis. Motion correction was performed using patch motion correction implemented in cryoSPARC. The contrast transfer function (CTF) was estimated using Patch Motion in cryoSPARC. Particles were picked using a blob picker (minimum particle diameter 100 Å, maximum particle diameter 150 Å). The particles of two datasets were extracted at a box size of 256 px and subjected to separate 2D classifications to identify good particles for *ab initio* model generation. *Ab initio* modelling with two volumes yielded one ‘good’ and one ‘junk’ map which were used in several rounds of heterorefinement. Heterorefinement resulted in a cleaned particle set (326,518) which was refined by homogeneous refinement using an automatically generated tight mask (dynamic mask near: 1 Å, dynamic mask far 3 Å, dynamic mask start resolution 4.5 Å). The particle set was further refined by non-uniform refinement with an automatically generated mask. The final 3D reconstruction yielded a resolution of 3.8 Å as estimated by the 0.143 cut-off criteria (Extended Data Fig. 3).

For truncated TSEN-pre-tRNA^Arg^tct, initial cryo-EM data analysis was performed in real-time using cryoSPARC live v. 3.3.1 (Structura Biotechnology Inc.). In the live application, 11,559 movies were collected with on-the-fly motion correction, CTF estimation, particle picking, and extraction. Initially, 9,961,856 particles were picked (minimum particle diameter 80 Å, maximum particle diameter 120 Å) and extracted at a 128 px box size. Particles were subjected to 2D classification in cryoSPARC and good particles were reextracted at a box size of 384 px. This data set was further polished by two rounds of 2D classification. Particle stacks were used to generate five ab initio models. After heterorefinement, one map represented 32.4% of the particle set (282,863) with similar appearance to the full-length TSEN/pre-tRNA complex. This set of particles was subjected to non-uniform refinement with an automatically generated mask. The resolution of the resulting map was estimated to 3.09 Å at the 0.143 FSC cut-off (Extended Data Fig. 3b).

### Reference models

The initial models for docking and manual adjustment in the cryo-EM maps were retrieved from the AlphFold Protein Structure Database^38,50^ with identifiers AF-Q8NCE0-F1 (TSEN2), AF-Q8WW01-F1 (TSEN15), AF-Q9BSV6-F1(TSEN34), and AF-Q7Z6J9-F1 (TSEN54). Other structures used in this study were retrieved from the protein data bank (PDB) with accession codes 6GJW (*Archaeglobus fulgidus* splicing endonuclease with BHB motif), 6Z9U (trunctated TSEN15–34 heterodimer), 3L0U (unmodified *Escherichia coli* tRNA^Phe^gaa), and 1EHZ (yeast tRNA^Phe^gaa).

### Model building, refinement, and analysis

Model building was initiated by docking the AlphaFold 3D structure predictions of the human proteins TSEN2, TSEN15, TSEN34, and TSEN54 ^38^ into the cryo-EM map followed by iterative rounds of automated flexible fitting of the models using Namdinator^51^ and manual adjustments in Coot^52^. Model building of pre-tRNA^Arg^tct, was guided using structures of unmodified *Escherichia coli* tRNA^Phe^gaa (PDB ID 3L0U) and yeast tRNA^Phe^gaa (PDB ID 1EHZ). Nucleotides 1-36 and 45-85 of pre-tRNA^Arg^tct, as well as residues 337-461 of TSEN2, residues 56-164 of TSEN15, residues 1-72 and 198-309 of TSEN34, and residues 30-174 and 446-520 of TSEN54, were finally build into the 3.1-Å cryo-EM density map. Real-space refinement of the model was carried out with Phenix^53,54^. MolProbity as a part of the Phenix validation tool was used for assessing the quality of the model^55,56^. Structural figures were rendered in PyMOL (The PyMOL Molecular Graphics System, Schrödinger), Chimera^57^, or ChimeraX^58^. Interface surface areas and molecular interactions were analyzed using PISA^59^. Data collection and model refinement statistics are summarized in Extended Data Table 1.

### All atom MD simulations

The simulation system for atomistic molecular dynamics simulations was set-up with CHARMM-GUI^60–62^ The protein was described by the AMBER14ff^63^, and the RNA with the OL3 force field^64^ and the TIP3P water model was employed^65^. The 3D structure of TSEN2 derived from the cryo-EM reconstruction was extended with the AlphaFold model^38^ and eventually comprised residues 1-143 and 217-459 of TSEN2. The model of pre-tRNA^Arg^tct, was completed by manually placing the missing nucleotides of the intron in Coot^52^ extending the A-I helix and a loop closure. Atomistic molecular dynamics simulations were run in GROMACS^66,67^. The system was neutralized and 150 mM KCl were added. After energy minimization, the system was relaxed for 125 ps with a 1 fs time step with position restraints. For the 1 μs production simulation the Nosé-Hoover thermostat with τt 1 ps and the Parinello-Rahman barostat with τp of 5 ps were used to maintain the temperature at 300 K and the pressure at 1 bar. A time step of 2 fs was used. The length of covalent bonds to hydrogens was constrained. The simulation system consisted of 330,000 atoms.

## Data availability

Cryo-EM density maps have been deposited in the Electron Microscopy Data Bank under accession numbers EMD-14925 (full-length TSEN/pre-tRNA^Tyr^gta complex) and EMD-14923 (truncated TSEN/pre-tRNA^Arg^tct complex). Atomic coordinates of truncated TSEN/pre-tRNA^Arg^tct complex were deposited to the Protein Data Bank (http://www.rcsb.org) under accession number 7ZRZ. MD simulation datasets were uploaded to the Zenodo Open Science repository (DOI 10.5281/zenodo.6513519). Uncropped versions of gels shown in Fig. 4e and Extended Data Fig. 1b,c are included as Supplementary Fig. 1. All other data needed to evaluate the conclusions in the paper are present in the paper and/or the supplementary materials. Relevant data are available from the corresponding author upon reasonable request.

## Code availability

All codes used in this study are freely available to academic users via the indicated resources.

## Acknowledgments

We thank Pascal Devant, Tim Julius Heinke, and Lina Sagert for biochemical characterization of TSEN and support in cell culture. We are grateful to Theresa Gewering and Arne Möller for initial single-particle EM analyses and Christian Kraft for technical support during cryo-EM data collection. Electron cryo-microscopy was carried out in the cryo-EM-facility of the Julius-Maximilians University Würzburg and the cryo-EM infrastructure of the Institute of Biochemistry and Collaborative Research Center 1507 (Z02 – high resolution cryo-EM platform). We thank Stefan Weitzer and Javier Martinez for critical reading of the manuscript. This study was supported by the Collaborative Research Center 902 ‘Molecular Principles of RNA-Based Regulation’ funded by the German Research Foundation. S.S. acknowledges a Boehringer Ingelheim Fonds fellowship. R.T. acknowledges funding by the European Research Council (ERC Advanced Grant No. 789121) and the German Research Foundation (Reinhart Koselleck Project No. TA157/12-1). S.T. acknowledges funding by the German Research Foundation (Grant No. TR 1711/1-7).

## Author contributions

S.S. produced and purified protein/RNA complexes, performed biochemical assays, and prepared cryo-EM grids. L.S. collected cryo-EM data and supported data processing carried out by S.S. and S.T.. S.S. and S.T. built the atomic model. L.S.S. performed all-atom molecular dynamics simulations. S.S. and S.T designed the experiments and wrote the initial draft of the manuscript with input from all authors. R.T. acquired funding. S.T. conceived the project, acquired funding, and supervised the work.

## Materials & correspondence

Correspondence and material requests should be addressed to the corresponding author.

## Tables

**Extended Data Table 1.**
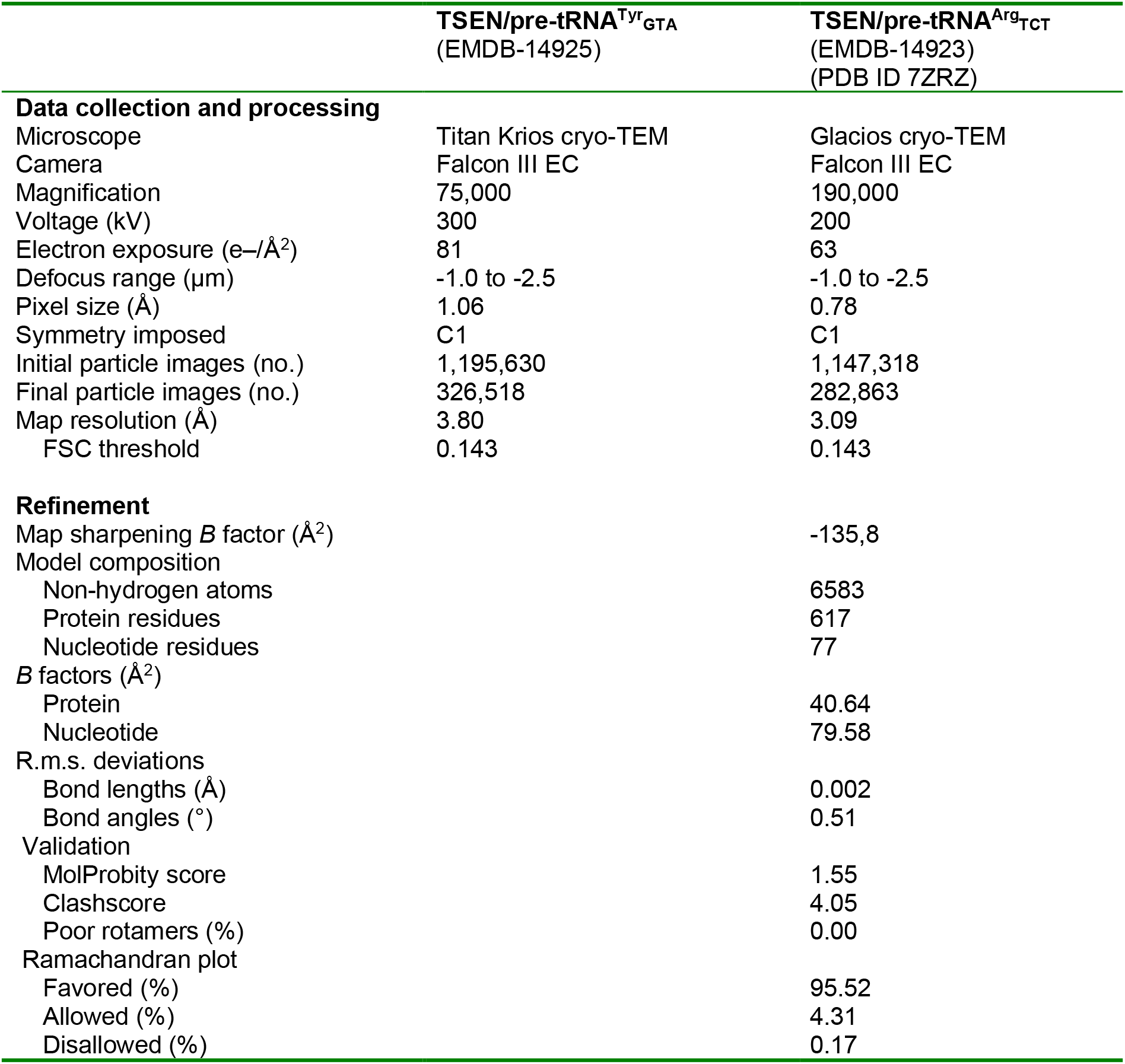
Cryo-EM data collection, refinement, and validation statistics for human TSEN maps and model.

|  | **TSEN/pre-tRNA<sup>Tyr</sup><sub>GTA</sub>**<br>(EMDB-14925) | **TSEN/pre-tRNA<sup>Arg</sup><sub>TCT</sub>**<br>(EMDB-14923)<br>(PDB ID 7ZRZ) |
| --- | --- | --- |
| **Data collection and processing** |  |  |
| Microscope | Titan Krios cryo-TEM | Glacios cryo-TEM |
| Camera | Falcon III EC | Falcon III EC |
| Magnification | 75,000 | 190,000 |
| Voltage (kV) | 300 | 200 |
| Electron exposure (e<sup>-</sup>/Å<sup>2</sup>) | 81 | 63 |
| Defocus range (μm) | -1.0 to -2.5 | -1.0 to -2.5 |
| Pixel size (Å) | 1.06 | 0.78 |
| Symmetry imposed | C1 | C1 |
| Initial particle images (no.) | 1,195,630 | 1,147,318 |
| Final particle images (no.) | 326,518 | 282,863 |
| Map resolution (Å) | 3.80 | 3.09 |
| FSC threshold | 0.143 | 0.143 |
| **Refinement** |  |  |
| Map sharpening *B* factor (Å<sup>2</sup>) |  | -135,8 |
| Model composition |  |  |
| Non-hydrogen atoms |  | 6583 |
| Protein residues |  | 617 |
| Nucleotide residues |  | 77 |
| *B* factors (Å<sup>2</sup>) |  |  |
| Protein |  | 40.64 |
| Nucleotide |  | 79.58 |
| R.m.s. deviations |  |  |
| Bond lengths (Å) |  | 0.002 |
| Bond angles (°) |  | 0.51 |
| Validation |  |  |
| MolProbity score |  | 1.55 |
| Clashscore |  | 4.05 |
| Poor rotamers (%) |  | 0.00 |
| Ramachandran plot |  |  |
| Favored (%) |  | 95.52 |
| Allowed (%) |  | 4.31 |
| Disallowed (%) |  | 0.17 |

**Supplementary Table 1.**
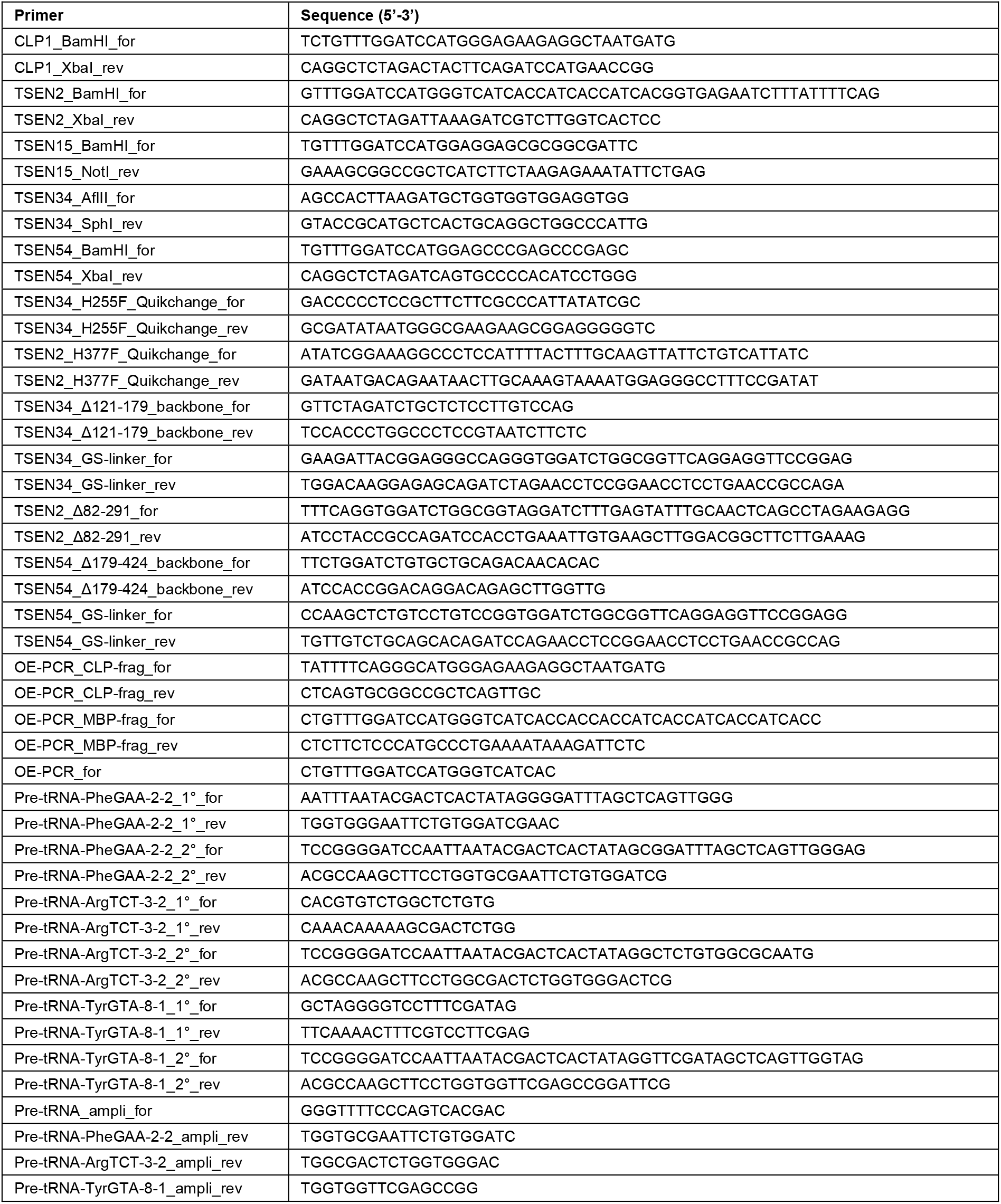
Primers used in this study.

## Figures

**Extended Data Fig. 1.**
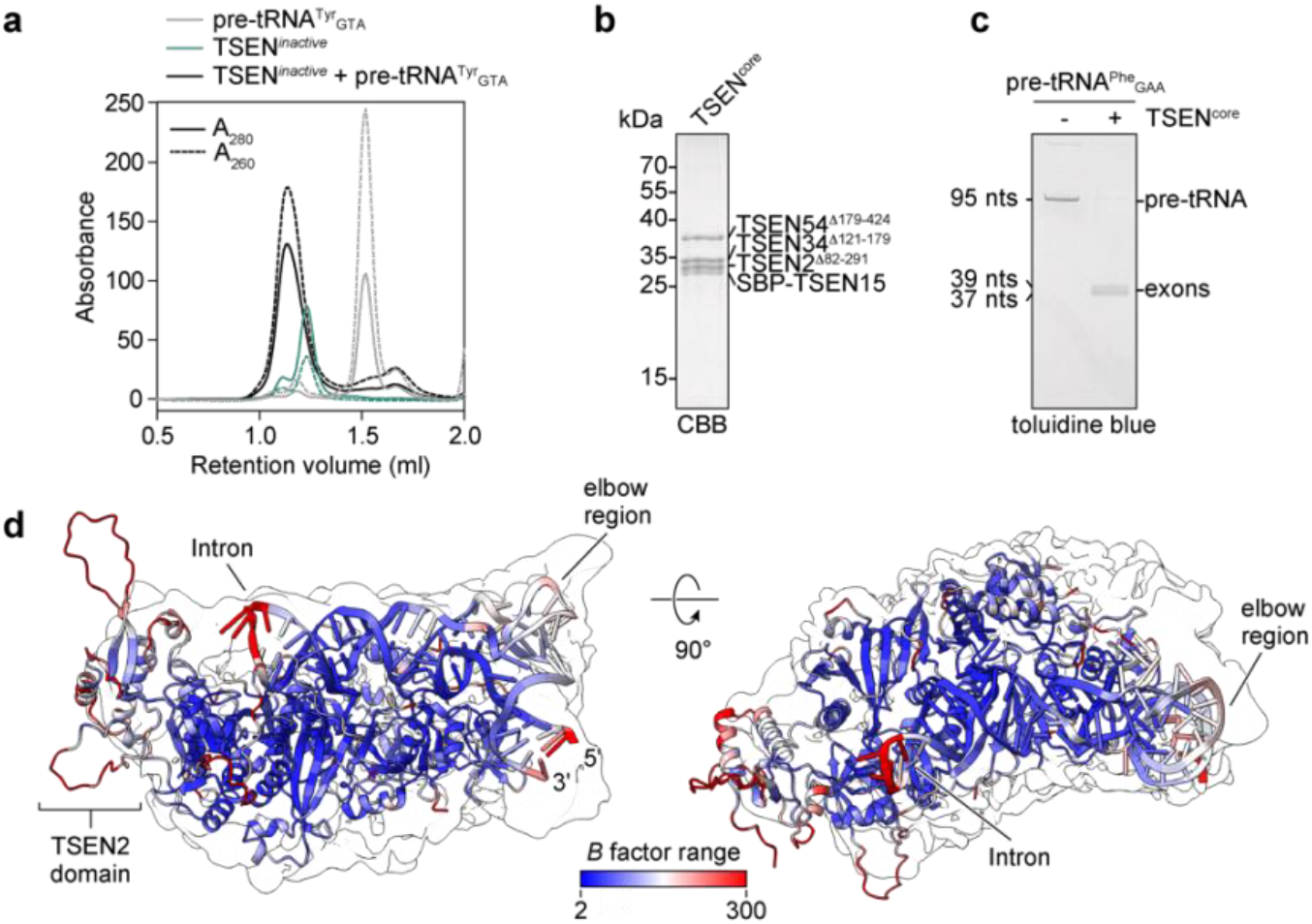
Biochemical and biophysical properties of TSEN/pre-tRNA complexes. **a**, Size exclusion chromatograms of TSEN/pre-tRNA^Tyr^gta complex and single components. Migration profile on a Superdex 200 Increase 3.2/300 column of TSEN/pre-tRNA^Tyr^gta complex (black), TSEN only (cyan) and pre-tRNA^Tyr^gta only (grey). **b**, SDS-PAGE analysis of truncated TSEN. **c**, Endonuclease activity assay with truncated TSEN. Yeast pre-tRNA^Phe^gaa was incubated without (left lane) and with TSEN (right lane) and analyzed by urea-PAGE (n = 2; independent biological replicates). **d**, Simulated flexibility of TSEN/pre-tRNA^Arg^tct. A hybrid model of truncated TSEN/pre-tRNA^Arg^tct with the central TSEN2 domain (residues 1-143 and 217-459) from the AlphaFold structure prediction^38^ and *in silico* modeled intron bases 37 to 44 was subjected to all-atom molecular dynamics simulations for assessment of structural flexibility. Color-coding is according to per residue *B* factor (Å^2^) with flexible regions in red and rigid regions in blue. The transparent local resolution filtered map of the complex is superimposed on the model.

**Extended Data Fig. 2.**
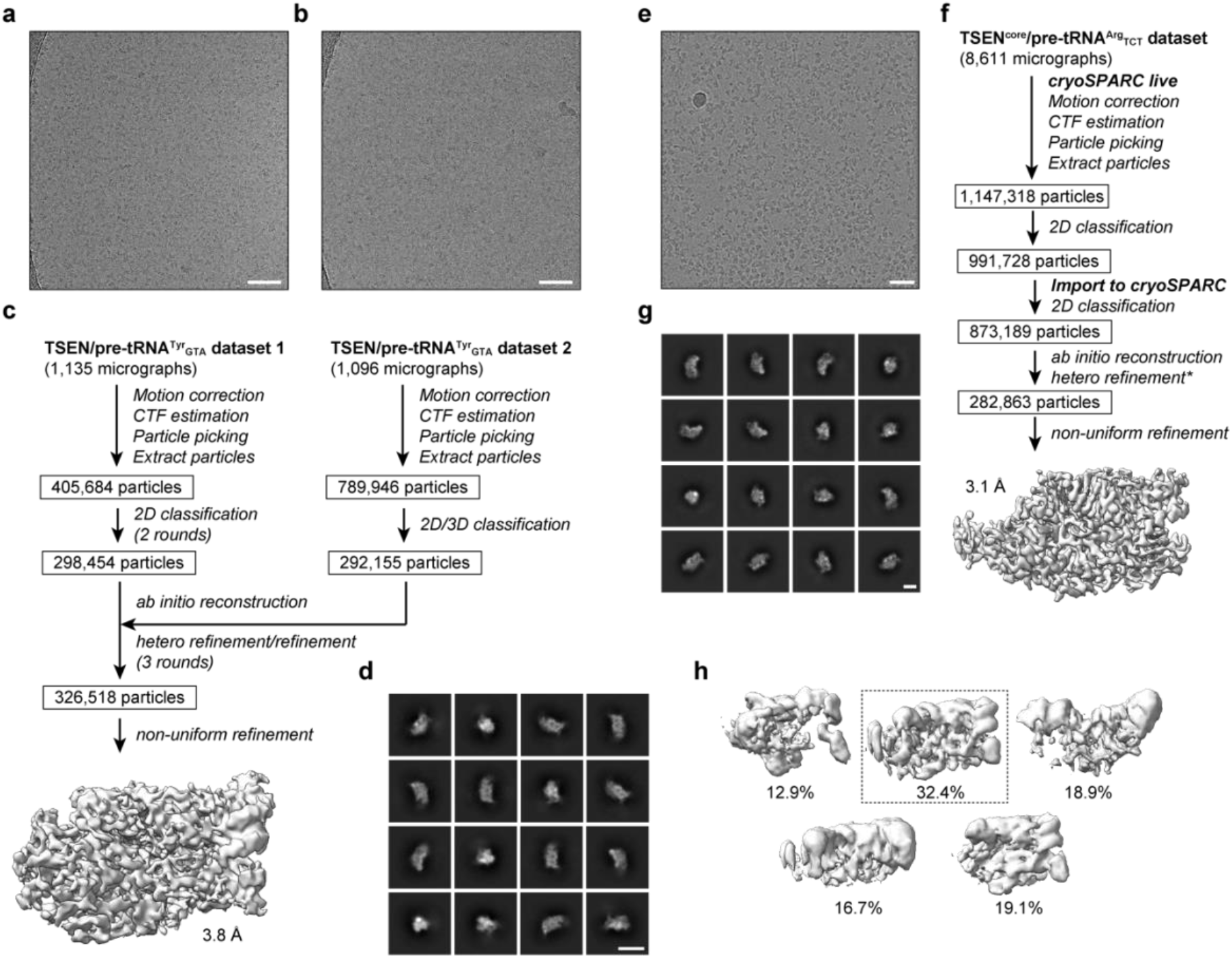
Cryo-EM data processing workflow of human TSEN (detailed in Methods). **a,b,e**, Representative micrographs of two TSEN/pre-tRNA^Tyr^gta datasets (**a,b**) and truncated TSEN/pre-tRNA^Arg^tct (**e**). White size markers indicate 50 nm. **c,f**, Workflow of cryo-EM data processing for TSEN/pre-tRNA^Tyr^gta (**c**) and truncated TSEN/pre-tRNA^Arg^tct (**f**) with final map and resolution. **d,g**, Representative 2D class images of TSEN/pre-tRNA^Tyr^gta (**d**) and TSEN/pre-tRNA^Arg^tct (**g**). White size markers indicate 10 nm. **h**, Heterorefined *ab initio* models of TSEN/pre-tRNA^Arg^tct. as shown in (**f**). Percentages indicate particle distribution among maps. The model in dotted box was further refined by non-uniform refinement.

**Extended Data Fig. 3.**
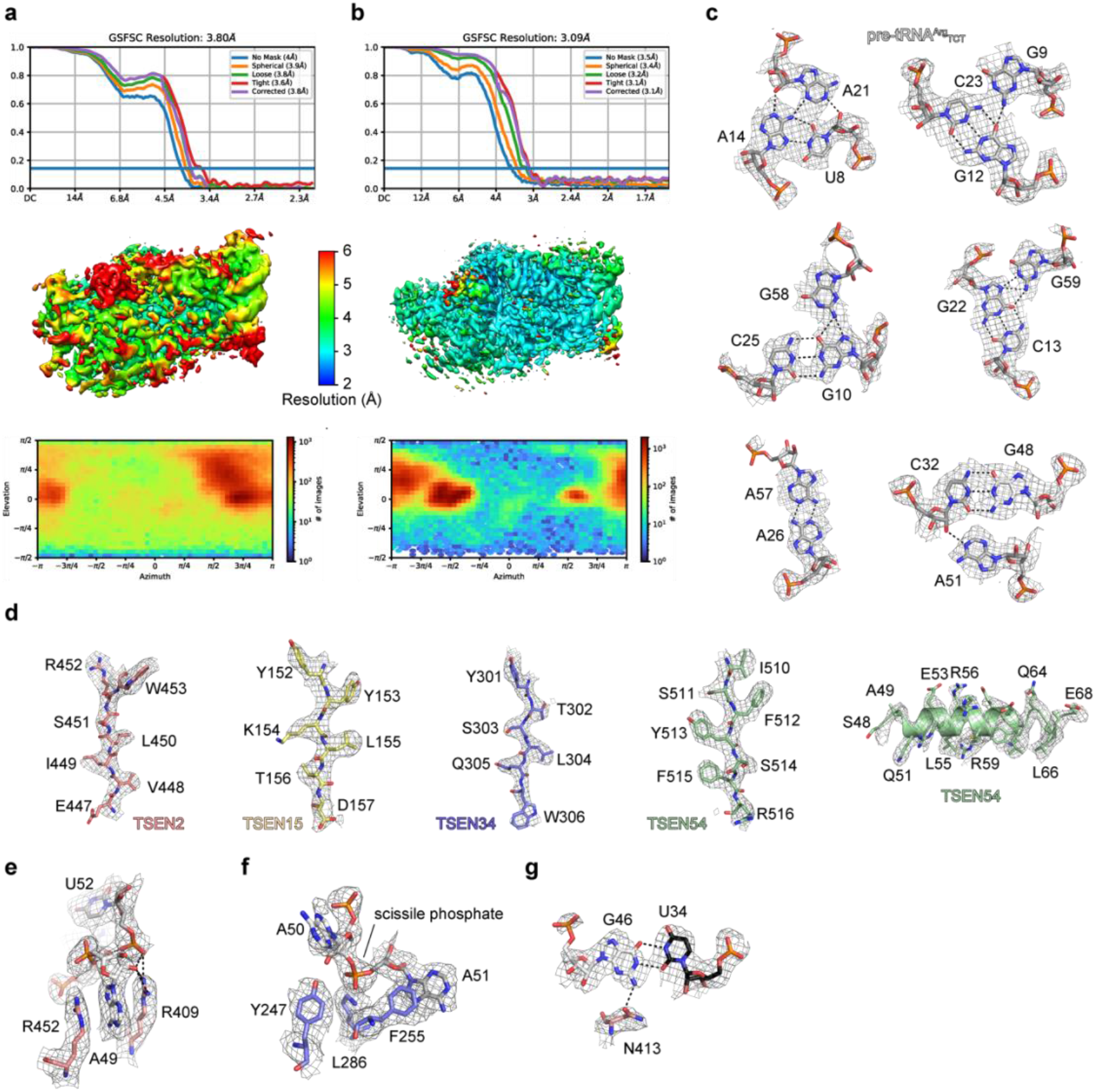
Quality of cryo-EM maps. **a,b**, Gold-standard Fourier shell correlation curves, local resolution maps, and Azimuth plots of TSEN/pre-tRNA^Tyr^gta (**a**) and TSEN/pre-tRNA^Arg^tct (**b**). Resolution values are derived from corrected curves without FSC-mask auto-tightening. The same color key was used for both local resolution maps. **c-f**, Representative cryo-EM densities enclosing the atomic models of pre-tRNA^Arg^tct secondary and tertiary structure base pairs (**c**), TSEN subunits (**d**), 3’ bulge cation-π sandwich (**e**), the 3’ splice site (**f**), and an interaction of TSEN2 N413 with the G-U wobble base pair of the A-I helix (**g**).

**Extended Data Fig. 4.**
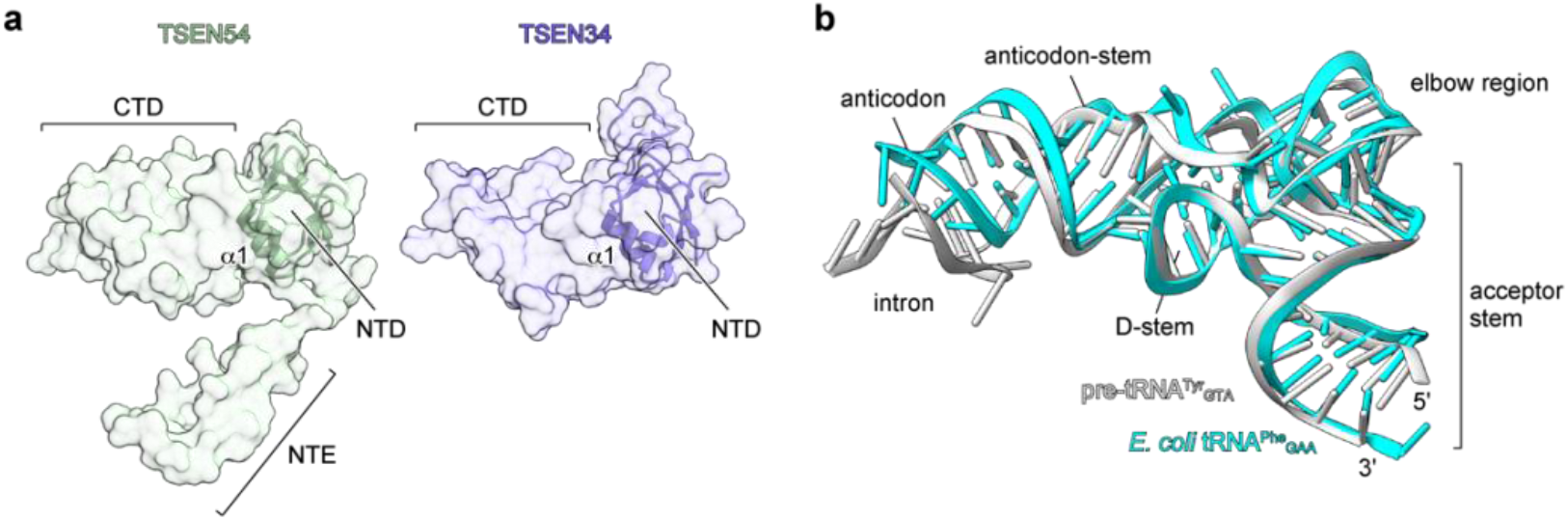
Structural aspects of TSEN/pre-tRNA complexes. **a**, Surface representation of TSEN54 and TSEN34 highlight their common N-terminal domains (NTD, cartoon representation). CTD – C-terminal domain, NTE – N-terminal extension. **b**, Superposition of *E. coli* tRNA^Phe^gaa (cyan; PDI ID 3L0U) onto human pre-tRNA^Arg^tct (grey, this work).

**Extended Data Fig. 5.**
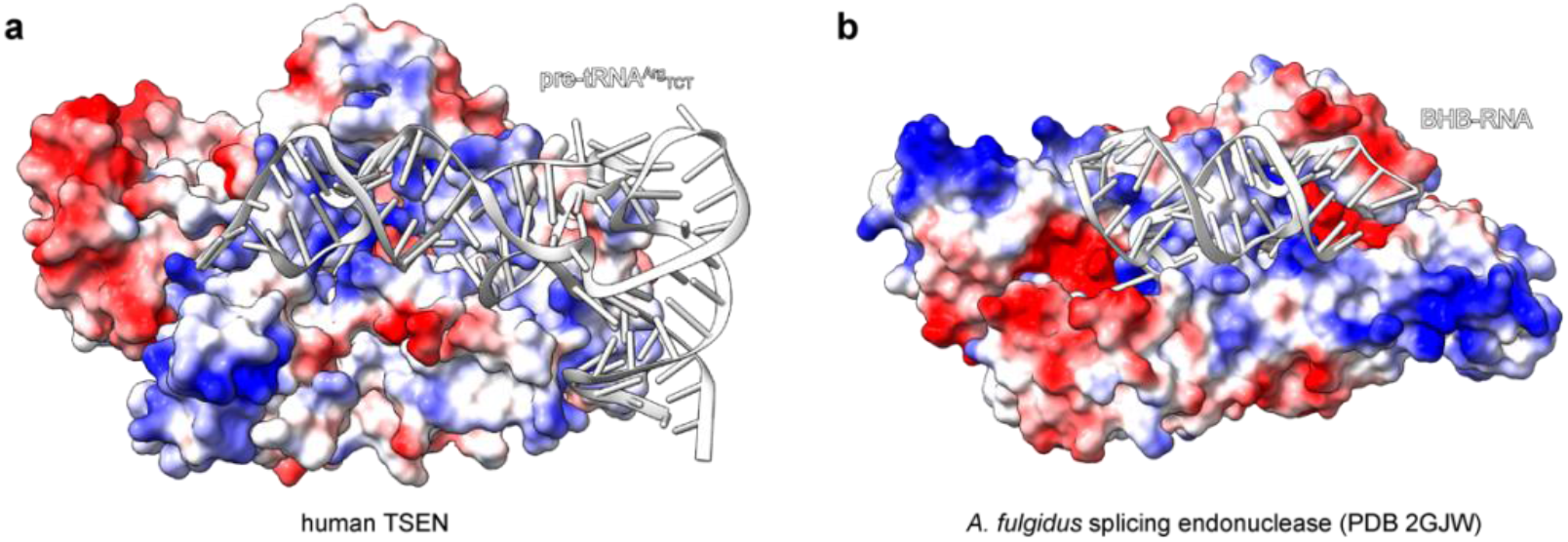
Electrostatic interactions of human and *A. fulgidus* splicing endonuclease. **a**, Electrostatic surface potential of human TSEN with pre-tRNA (grey). **b**, Electrostatic surface potential of *Archaeoglobus fulgidus* endonuclease (PDB ID 2GJW) with BHB-RNA (grey). For comparison, the Archaeal endonuclease structure was superimposed with the cryo-EM model of TSEN-pre-tRNA^Arg^tct.

**Extended Data Fig. 6.**
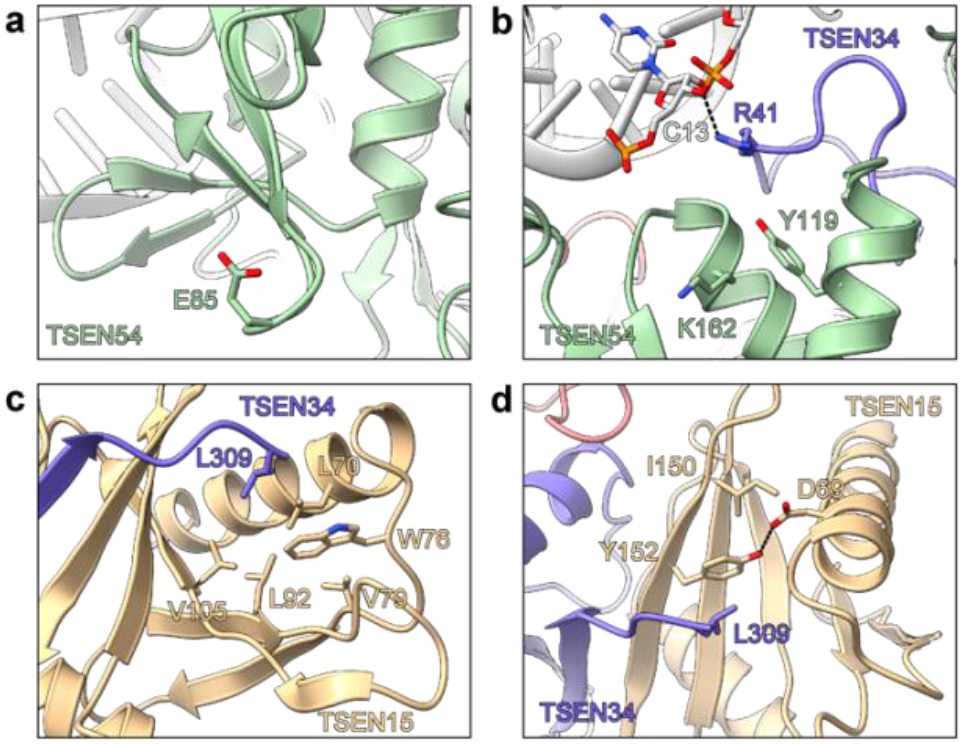
Molecular details of TSEN interactions in regions of PCH mutation hotspots. **a**, TSEN54 (green) S85 environment in proximity to pre-tRNA acceptor stem (grey). **b**, Environment of the TSEN54 Y119D mutation potentially impacting pre-tRNA (grey) recognition. **c**, Environment of the TSEN15 (yellow) W76 environment. TSEN34 is shown in slate blue. **d**, TSEN15 Y152 environment.

**Supplementary Fig. 1.**
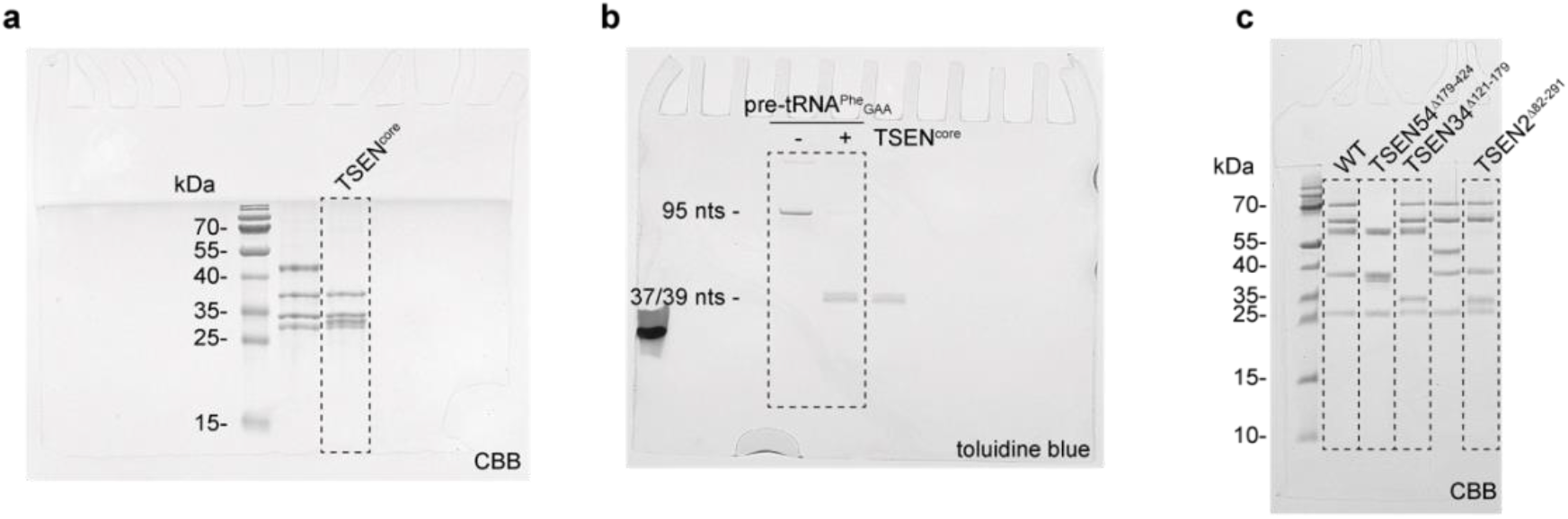
Source images for all data obtained by gel electrophoresis. **a**, Related to extended data figure 1b. **b**, Related to extended data figure 1c. **c**, Related to figure 4e.

## References

1 Paushkin, S. V., Patel, M., Furia, B. S., Peltz, S. W. & Trotta, C. R. Identification of a human endonuclease complex reveals a link between tRNA splicing and pre-mRNA 3’ end formation. Cell 117, 311–321, (2004).

2 Budde, B. S. et al. tRNA splicing endonuclease mutations cause pontocerebellar hypoplasia. Nat Genet 40, 1113–1118, (2008).

3 Namavar, Y. et al. Clinical, neuroradiological and genetic findings in pontocerebellar hypoplasia. Brain 134, 143–156, (2011).

4 Bierhals, T., Korenke, G. C., Uyanik, G. & Kutsche, K. Pontocerebellar hypoplasia type 2 and TSEN2: review of the literature and two novel mutations. Eur J Med Genet 56, 325–330, (2013).

5 Alazami, A. M. et al. Accelerating novel candidate gene discovery in neurogenetic disorders via whole-exome sequencing of prescreened multiplex consanguineous families. Cell Rep 10, 148–161, (2015).

6 Breuss, M. W. et al. Autosomal-Recessive Mutations in the tRNA Splicing Endonuclease Subunit TSEN15 Cause Pontocerebellar Hypoplasia and Progressive Microcephaly. Am J Hum Genet 99, 785, (2016).

7 Karaca, E. et al. Human CLP1 mutations alter tRNA biogenesis, affecting both peripheral and central nervous system function. Cell 157, 636–650, (2014).

8 Schaffer, A. E. et al. CLP1 founder mutation links tRNA splicing and maturation to cerebellar development and neurodegeneration. Cell 157, 651–663, (2014).

9 Li, H., Trotta, C. R. & Abelson, J. Crystal structure and evolution of a transfer RNA splicing enzyme. Science 280, 279–284, (1998).

10 Sekulovski, S. et al. Assembly defects of human tRNA splicing endonuclease contribute to impaired pre-tRNA processing in pontocerebellar hypoplasia. Nat Commun 12, 5610, (2021).

11 Schimmel, P. The emerging complexity of the tRNA world: mammalian tRNAs beyond protein synthesis. Nat Rev Mol Cell Biol 19, 45–58, (2018).

12 Chan, P. P. & Lowe, T. M. GtRNAdb 2.0: an expanded database of transfer RNA genes identified in complete and draft genomes. Nucleic Acids Res 44, D184–189, (2016).

13 Popow, J. et al. HSPC117 is the essential subunit of a human tRNA splicing ligase complex. Science 331, 760–764, (2011).

14 Rauhut, R., Green, P. R. & Abelson, J. Yeast tRNA-splicing endonuclease is a heterotrimeric enzyme. J Biol Chem 265, 18180–18184, (1990).

15 Song, J. & Markley, J. L. Three-dimensional structure determined for a subunit of human tRNA splicing endonuclease (Sen15) reveals a novel dimeric fold. J Mol Biol 366, 155–164, (2007).

16 Tocchini-Valentini, G. D., Fruscoloni, P. & Tocchini-Valentini, G. P. Structure, function, and evolution of the tRNA endonucleases of Archaea: an example of subfunctionalization. Proc Natl Acad Sci U S A 102, 8933–8938, (2005).

17 Hirata, A. et al. X-ray structure of the fourth type of archaeal tRNA splicing endonuclease: insights into the evolution of a novel three-unit composition and a unique loop involved in broad substrate specificity. Nucleic Acids Res 40, 10554–10566, (2012).

18 Trotta, C. R. et al. The yeast tRNA splicing endonuclease: a tetrameric enzyme with two active site subunits homologous to the archaeal tRNA endonucleases. Cell 89, 849–858, (1997).

19 Hayne, C. K., Schmidt, C. A., Haque, M. I., Matera, A. G. & Stanley, R. E. Reconstitution of the human tRNA splicing endonuclease complex: insight into the regulation of pre-tRNA cleavage. Nucleic Acids Res, (2020).

20 Lee, M. C. & Knapp, G. Transfer RNA splicing in Saccharomyces cerevisiae. Secondary and tertiary structures of the substrates. J Biol Chem 260, 3108–3115, (1985).

21 Swerdlow, H. & Guthrie, C. Structure of intron-containing tRNA precursors. Analysis of solution conformation using chemical and enzymatic probes. J Biol Chem 259, 5197–5207, (1984).

22 Thompson, L. D. & Daniels, C. J. Recognition of exon-intron boundaries by the Halobacterium volcanii tRNA intron endonuclease. J Biol Chem 265, 18104–18111, (1990).

23 Reyes, V. M. & Abelson, J. Substrate recognition and splice site determination in yeast tRNA splicing. Cell 55, 719–730, (1988).

24 Greer, C. L., Soll, D. & Willis, I. Substrate recognition and identification of splice sites by the tRNA-splicing endonuclease and ligase from Saccharomyces cerevisiae. Mol Cell Biol 7, 76–84, (1987).

25 Baldi, M. I., Mattoccia, E., Bufardeci, E., Fabbri, S. & Tocchini-Valentini, G. P. Participation of the intron in the reaction catalyzed by the Xenopus tRNA splicing endonuclease. Science 255, 1404–1408, (1992).

26 Di Nicola Negri, E. et al. The eucaryal tRNA splicing endonuclease recognizes a tripartite set of RNA elements. Cell 89, 859–866, (1997).

27 Schmidt, C. A., Giusto, J. D., Bao, A., Hopper, A. K. & Matera, A. G. Molecular determinants of metazoan tricRNA biogenesis. Nucleic Acids Res 47, 6452–6465, (2019).

28 Xue, S., Calvin, K. & Li, H. RNA recognition and cleavage by a splicing endonuclease. Science 312, 906–910, (2006).

29 Trotta, C. R., Paushkin, S. V., Patel, M., Li, H. & Peltz, S. W. Cleavage of pre-tRNAs by the splicing endonuclease requires a composite active site. Nature 441, 375–377, (2006).

30 Weitzer, S. & Martinez, J. The human RNA kinase hClp1 is active on 3’ transfer RNA exons and short interfering RNAs. Nature 447, 222–226, (2007).

31 Hanada, T. et al. CLP1 links tRNA metabolism to progressive motor-neuron loss. Nature 495, 474–480, (2013).

32 Monaghan, C. E., Adamson, S. I., Kapur, M., Chuang, J. H. & Ackerman, S. L. The Clp1 R140H mutation alters tRNA metabolism and mRNA 3’ processing in mouse models of pontocerebellar hypoplasia. Proc Natl Acad Sci U S A 118, (2021).

33 Schmidt, C. A. et al. Mutations in Drosophila tRNA processing factors cause phenotypes similar to Pontocerebellar Hypoplasia. Biol Open 11, (2022).

34 van Dijk, T., Baas, F., Barth, P. G. & Poll-The, B. T. What’s new in pontocerebellar hypoplasia? An update on genes and subtypes. Orphanet J Rare Dis 13, 92, (2018).

35 Schmidt, C. A. & Matera, A. G. tRNA introns: Presence, processing, and purpose. Wiley Interdiscip Rev RNA 11, e1583, (2020).

36 Weitzer, S., Hanada, T., Penninger, J. M. & Martinez, J. CLP1 as a novel player in linking tRNA splicing to neurodegenerative disorders. Wiley Interdiscip Rev RNA 6, 47–63, (2015).

37 Gogakos, T. et al. Characterizing expression and processing of precursor and mature human tRNAs by hydro-tRNAseq and PAR-CLIP. Cell Rep 20, 1463–1475, (2017).

38 Jumper, J. et al. Highly accurate protein structure prediction with AlphaFold. Nature 596, 583–589, (2021).

39 Byrne, R. T., Konevega, A. L., Rodnina, M. V. & Antson, A. A. The crystal structure of unmodified tRNAPhe from Escherichia coli. Nucleic Acids Res 38, 4154–4162, (2010).

40 Bufardeci, E., Fabbri, S., Baldi, M. I., Mattoccia, E. & Tocchini-Valentini, G. P. In vitro genetic analysis of the structural features of the pre-tRNA required for determination of the 3’ splice site in the intron excision reaction. EMBO J 12, 4697–4704, (1993).

41 Belfort, M. & Weiner, A. Another bridge between kingdoms: tRNA splicing in archaea and eukaryotes. Cell 89, 1003–1006, (1997).

42 Hurtig, J. E. et al. Comparative parallel analysis of RNA ends identifies mRNA substrates of a tRNA splicing endonuclease-initiated mRNA decay pathway. Proc Natl Acad Sci U S A 118, (2021).

43 de Vries, H. et al. Human pre-mRNA cleavage factor II(m) contains homologs of yeast proteins and bridges two other cleavage factors. EMBO J 19, 5895–5904, (2000).

44 Noble, C. G., Beuth, B. & Taylor, I. A. Structure of a nucleotide-bound Clp1-Pcf11 polyadenylation factor. Nucleic Acids Res 35, 87–99, (2007).

45 Miao, F. & Abelson, J. Yeast tRNA-splicing endonuclease cleaves precursor tRNA in a random pathway. J Biol Chem 268, 672–677, (1993).

46 Li, M. Z. & Elledge, S. J. Harnessing homologous recombination in vitro to generate recombinant DNA via SLIC. Nat Methods 4, 251–256, (2007).

47 Reyes, V. M. & Abelson, J. A synthetic substrate for tRNA splicing. Anal Biochem 166, 90–106, (1987).

48 Easton, L. E., Shibata, Y. & Lukavsky, P. J. Rapid, nondenaturing RNA purification using weak anion-exchange fast performance liquid chromatography. RNA 16, 647–653, (2010).

49 Punjani, A., Rubinstein, J. L., Fleet, D. J. & Brubaker, M. A. cryoSPARC: algorithms for rapid unsupervised cryo-EM structure determination. Nat Methods 14, 290–296, (2017).

50 Varadi, M. et al. AlphaFold Protein Structure Database: massively expanding the structural coverage of protein-sequence space with high-accuracy models. Nucleic Acids Res 50, D439–D444, (2022).

51 Kidmose, R. T. et al. Namdinator - automatic molecular dynamics flexible fitting of structural models into cryo-EM and crystallography experimental maps. IUCrJ 6, 526–531, (2019).

52 Emsley, P., Lohkamp, B., Scott, W. G. & Cowtan, K. Features and development of Coot. Acta Crystallogr D Biol Crystallogr 66, 486–501, (2010).

53 Adams, P. D. et al. PHENIX: a comprehensive Python-based system for macromolecular structure solution. Acta Crystallogr D Biol Crystallogr 66, 213–221, (2010).

54 Liebschner, D. et al. Macromolecular structure determination using X-rays, neutrons and electrons: recent developments in Phenix. Acta Crystallogr D Struct Biol 75, 861–877, (2019).

55 Chen, V. B. et al. MolProbity: all-atom structure validation for macromolecular crystallography. Acta Crystallogr D Biol Crystallogr 66, 12–21, (2010).

56 Williams, C. J. et al. MolProbity: More and better reference data for improved all-atom structure validation. Protein Sci 27, 293–315, (2018).

57 Pettersen, E. F. et al. UCSF Chimera--a visualization system for exploratory research and analysis. J Comput Chem 25, 1605–1612, (2004).

58 Pettersen, E. F. et al. UCSF ChimeraX: Structure visualization for researchers, educators, and developers. Protein Sci 30, 70–82, (2021).

59 Krissinel, E. & Henrick, K. Inference of macromolecular assemblies from crystalline state. J Mol Biol 372, 774–797, (2007).

60 Jo, S., Kim, T., Iyer, V. G. & Im, W. CHARMM-GUI: a web-based graphical user interface for CHARMM. J Comput Chem 29, 1859–1865, (2008).

61 Brooks, B. R. et al. CHARMM: the biomolecular simulation program. J Comput Chem 30, 1545–1614, (2009).

62 Lee, J. et al. CHARMM-GUI Input Generator for NAMD, GROMACS, AMBER, OpenMM, and CHARMM/OpenMM Simulations Using the CHARMM36 Additive Force Field. J Chem Theory Comput 12, 405–413, (2016).

63 Maier, J. A. et al. ff14SB: Improving the Accuracy of Protein Side Chain and Backbone Parameters from ff99SB. J Chem Theory Comput 11, 3696–3713, (2015).

64 Zgarbova, M. et al. Refinement of the Cornell et al. Nucleic Acids Force Field Based on Reference Quantum Chemical Calculations of Glycosidic Torsion Profiles. J Chem Theory Comput 7, 2886–2902, (2011).

65 Jorgensen, W. L., Chandrasekhar, J., Madura, J. D., Impey, R. W. & Klein, M. L. Comparison of simple potential functions for simulating liquid water. Journal Chemical Physics 79, 926–935, (1983).

66 Pronk, S. et al. GROMACS 4.5: a high-throughput and highly parallel open source molecular simulation toolkit. Bioinformatics 29, 845–854, (2013).

67 Abraham, M. J. et al. GROMACS: High performance molecular simulations through multi-level parallelism from laptops to supercomputers. SoftwareX 1–2, 19–25, (2015).

